# *Caulobacter crescentus* RNase E condensation contributes to autoregulation and fitness

**DOI:** 10.1101/2023.12.15.571756

**Authors:** Vidhyadhar Nandana, Nadra Al-Husini, Arti Vaishnav, Kulathungage H. Dilrangi, Jared M. Schrader

## Abstract

RNase E is the most common RNA decay nuclease in bacteria, setting the global mRNA decay rate and scaffolding formation of the RNA degradosome complex and BR-bodies. To properly set the global mRNA decay rate, RNase E from *Escherichia coli* and neighboring γ-proteobacteria were found to autoregulate RNase E levels via the decay of its mRNA’s 5’ UTR. While the 5’ UTR is absent from other groups of bacteria in the Rfam database, we identified that the α-proteobacterium *Caulobacter crescentus* RNase E contains a similar 5’ UTR structure that promotes RNase E autoregulation. In both bacteria, the C-terminal IDR of RNase E is required for proper autoregulation to occur, and this IDR is also necessary and sufficient for RNase E to phase-separate, generating BR-bodies. Using *in vitro* purified RNase E, we find that the IDR’s ability to promote phase-separation correlates with enhanced 5’ UTR cleavage, suggesting that phase-separation of RNase E with the 5’ UTR enhances autoregulation. Finally, using growth competition experiments we find that a strain capable of autoregulation rapidly outcompetes a strain with a 5’ UTR mutation that cannot autoregulate, suggesting autoregulation promotes optimal cellular fitness.

## Introduction

In bacteria, mRNA decay is typically controlled by the protein RNase E (1–5). RNase E is an endonuclease that performs the rate-limiting step of RNA cleavage, setting the global rate of mRNA decay (6, 7). To do this, the levels of RNase E in *Escherichia coli* are carefully controlled through autoregulation activity whereby the protein binds to the 5’ UTR in its own mRNA and facilitates its cleavage (8, 9). Importantly, this mechanism of autoregulation allows the cell to adjust its mRNA decay demands to changes in mRNA abundance (10). While the 5’ UTR is well conserved among γ-proteobacteria (11), it is currently not annotated outside this clade of bacteria in Rfam (12). In *Caulobacter crescentus*, an α-proteobacterium, it was observed that RNase E can also autoregulate its expression (13), however, the 5’ UTR secondary structure analysis and functional impact on autoregulation were not explicitly investigated. In E. *coli* and *C. crescentus*, it was found that the intrinsically disordered C-terminal domain of RNase E is necessary for autoregulation (13, 14). Interestingly, this C-terminal region of RNase E is also necessary and sufficient for the formation of BR-bodies, phase-separated biomolecular condensates that promote mRNA decay activity (7). However, the role of BR-bodies in the process of RNase E autoregulation has not been directly tested. Finally, while RNase E appears to be essential in *C. crescentus* (13), it has not been thoroughly tested as to how autoregulation of RNase E contributes to cellular fitness.

To examine the role of RNase E autoregulation in *C. crescentus*, we show that under- or overexpression of RNase E led to significant reductions in cell growth. We find that like the γ-proteobacteria, the α-proteobacteria likely also utilize a similar 5’ UTR structure that is necessary for RNase E autoregulation. By performing a growth competition experiment, a mutant whose 5’ UTR was replaced by a synthetic 5’ UTR, was rapidly outcompeted, suggesting that even mild overexpression of RNase E lowers fitness. Finally, using an *in vitro* purified system, we find that in the presence of *C. crescentus* RNase E condensates, 5’ UTR cleavage is stimulated, suggesting phase-separation of RNase E together with the 5’ UTR promotes RNase E cleavage leading to autoregulation.

## Results

### RNase E depletion or overexpression leads to loss of fitness

To determine the essentiality of the major mRNA endonuclease RNase E, we used a depletion strain (JS8) where the promoter of RNase E was replaced with a xylose inducible promoter (Fig 1A). While a previous Tn-seq study found it was a high fitness cost gene in rich media, insertions were only found in the intrinsically disordered CTD and were absent in the catalytic NTD, suggesting its activity may be essential for growth(15). In liquid culture, we found that the depletion strain had somewhat attenuated growth compared to a control strain in the presence of xylose, but the observed growth rate was slower in the absence of xylose (Fig 1A). The deceleration of growth likely arises from the slow depletion of the RNase E protein by cell division which takes approximately 4 hours (Fig 1A). After 8-hours of growth without xylose, the depletion strain shows an ∼4 log reduction in the number of colonies formed compared to the control, suggesting the gene is indeed critical for growth (Fig 1B). Of note, the RNase E depletion strain colonies that grew in the absence of xylose were heterogenous in size and the larger ones no longer required xylose to grow in liquid cultures (data not shown), suggesting they accumulated mutations that allow constitutive expression of RNase E in the absence of xylose. Additionally, our attempts to make a clean deletion of the RNase E gene failed in our hands (data not shown), suggesting the RNase E gene is likely essential, or at a minimum provides very high fitness cost in *C. crescentus*. Altogether, this suggests that depletion of RNase E leads to a strong reduction in cell growth.

**Figure 1.**
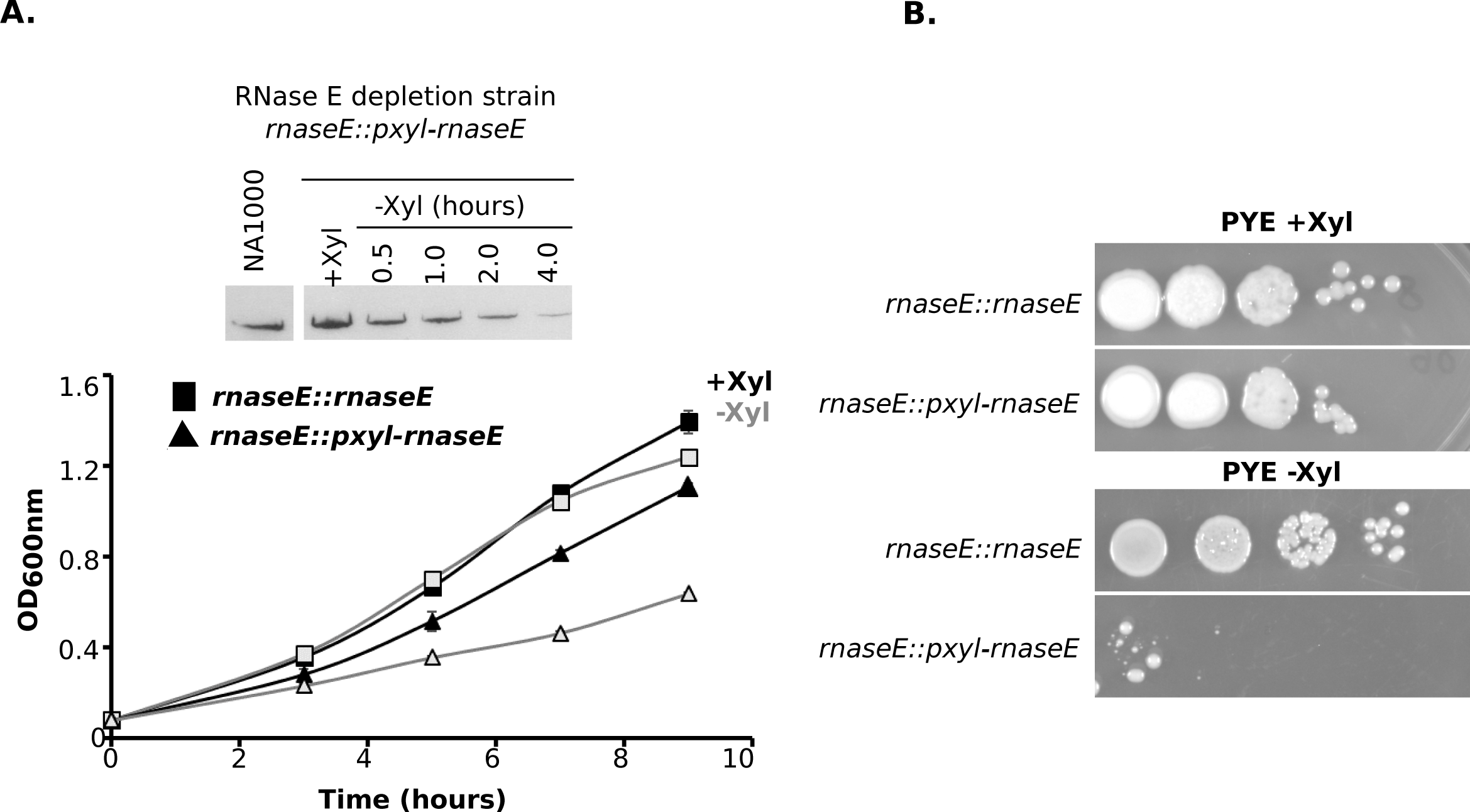
Depletion of RNase E leads to slow growth and loss in viability. A.) Growth curves of depletion strains of RNase E (triangles) and empty vector controls (squares) in the presence of xylose (black) or the absence of xylose (light grey). All cells were grown in PYE with kanamycin. B.) Colony forming unit assay to determine cell viability in the Depletion strains. Depletion strains were grown in PYE/Kan in the presence of xylose (black) or absence of xylose (light grey) for 8 hours, then spotted on PYE/Kan/Xyl plates. EV = Empty vector, RNE=RNase E. C.) Colony forming unit assay to determine cell viability in the RNase E replacement strains. Replacement strains contain a xylose inducible copy of RNase E at the *rne* locus, and mutant RNase E variants at the *vanA* locus. RNase E variants expressed from the *vanA* locus are indicated to the left. FL = full length, ΔEG = catalytic domain deletion, ASM = active site mutant.

Next, we explored the functional consequences of artificial overexpression of RNase E. We generated a pBX multicopy plasmid containing the RNase E-YFP construct (JS89) (Fig 2A). Cells harboring an empty vector showed little difference in growth in the presence or absence of xylose (Fig 2A). In the strain harboring the plasmid with RNase E-YFP, we found that growth was significantly slower in the absence of xylose, likely due to leaky levels of expression that surpassed that of the wild-type cells (Fig 2A), however, growth was halted when the xylose inducer was added (Fig 2A). Additionally, we added the xylose inducer to cells as they approached an OD600 of 0.3 in the middle of log-phase instead of at OD600 of 0.05 and found that the growth rate was maintained similar to the uninduced cells for approximately 2-hours, but then showed a significant reduction after 3-hours (Fig 2A). The maximum induction of RNase E protein appeared to occur after 3-hours of induction with xylose (Fig S1), suggesting that the reduction in growth observed at 3-hours is due to its peak in protein accumulation. Of note, we observed that when blotted using anti-RNase E antibodies, the chromosomal copy of RNase E’s expression was no longer detected after RNase E overexpression, suggesting that it was shut off in response to plasmid-based expression (Fig 2A). To examine if the cells were losing viability after RNase E overexpression, we also performed dilution assays, plated the dilutions on solid media, and assayed for colony formation (Fig 2B). Here, induction of RNase E led to a complete loss in colony formation upon RNase E overexpression (Fig 2B). In conclusion, when RNase E is overexpressed strongly above wild-type levels, *C. crescentus* has a strong reduced growth rate and reduction in cell viability.

**Figure 2.**
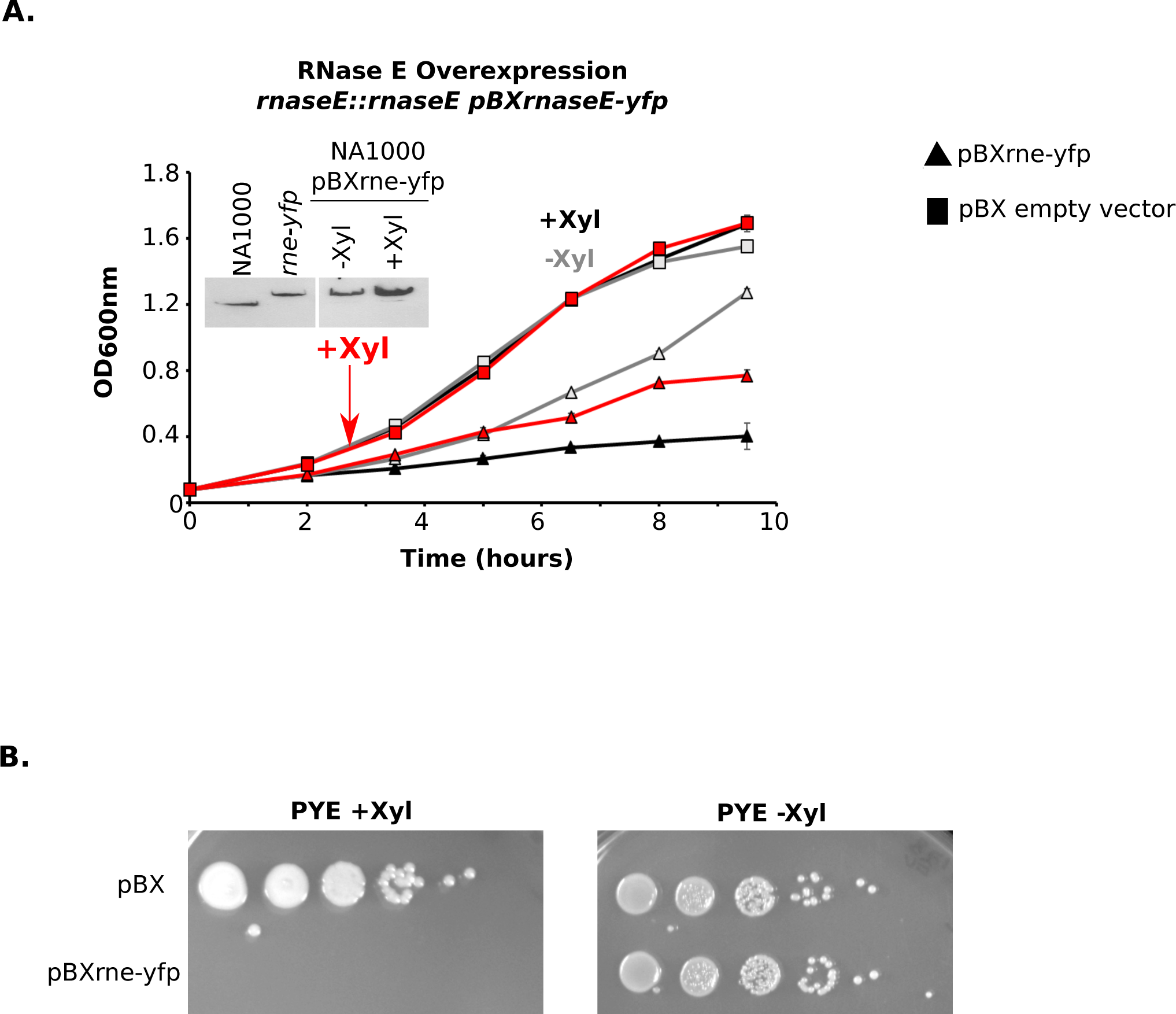
Overexpression of RNase E leads to growth arrest. A.) Growth curves of overexpression strains of RNase E (triangles), and empty vector controls (squares) grown in the presence of xylose (black) or the absence of xylose (light grey). Cells induced with xylose after the 2-hour timepoint are colored red. All cells were grown in PYE with kanamycin. B.) Colony Forming unit assay to determine cell viability in the overexpression strains harboring pBX vectors with the insert on the left.

### RNase E autoregulation ensures optimal cell fitness

RNase E is known to autoregulate its own levels in *E. coli* and other γ-proteobacteria (8, 9, 11, 13, 16) and we previously observed autoregulation of *C. crescentus* RNase E when expressed from the *vanA* locus on the chromosome (13). Here, we noticed that when cells harbored an extra copy of RNase E-YFP on the pBX plasmid, chromosomal RNase E levels from the *rne* locus were no longer detected (Fig 2A). Further, when a second copy of RNase E-YFP with a plasmid derived 5’ UTR was expressed artificially from the vanillate promoter, we observed that the relative abundance of the wild-type RNase E protein was reduced by a corresponding amount (Fig 3A). To further examine the properties of RNase E autoregulation, we induced copy of RNaseE-YFP from the *vanA* locus at different levels of vanillate and found that the more RNase E-YFP produced, the stronger the inhibition of native RNase E expression (Fig 3A). In *E. coli*, the 5’ UTR of RNase E has an RNA structural element which is recognized by RNase E that is necessary for its autoregulation (8). To test whether the *C. crescentus* RNase E gene’s 5’ UTR is responsible for autoregulation, we generated a strain in which an active site mutant of RNase E that is unable to cleave RNA (7) was expressed from the chromosome (Fig 3A). This version failed to shut down expression of the native RNase E gene (13), suggesting that RNase E activity is required for autoregulation. Additionally, by using the strain in which the native RNase E’s 5’ UTR was replaced by a plasmid derived ribosome binding site, we also observed a failure to autoregulate *rne* expression (Fig 3A). Altogether, this suggests that RNase E activity on its 5’ UTR is necessary for autoregulation.

**Figure 3.**
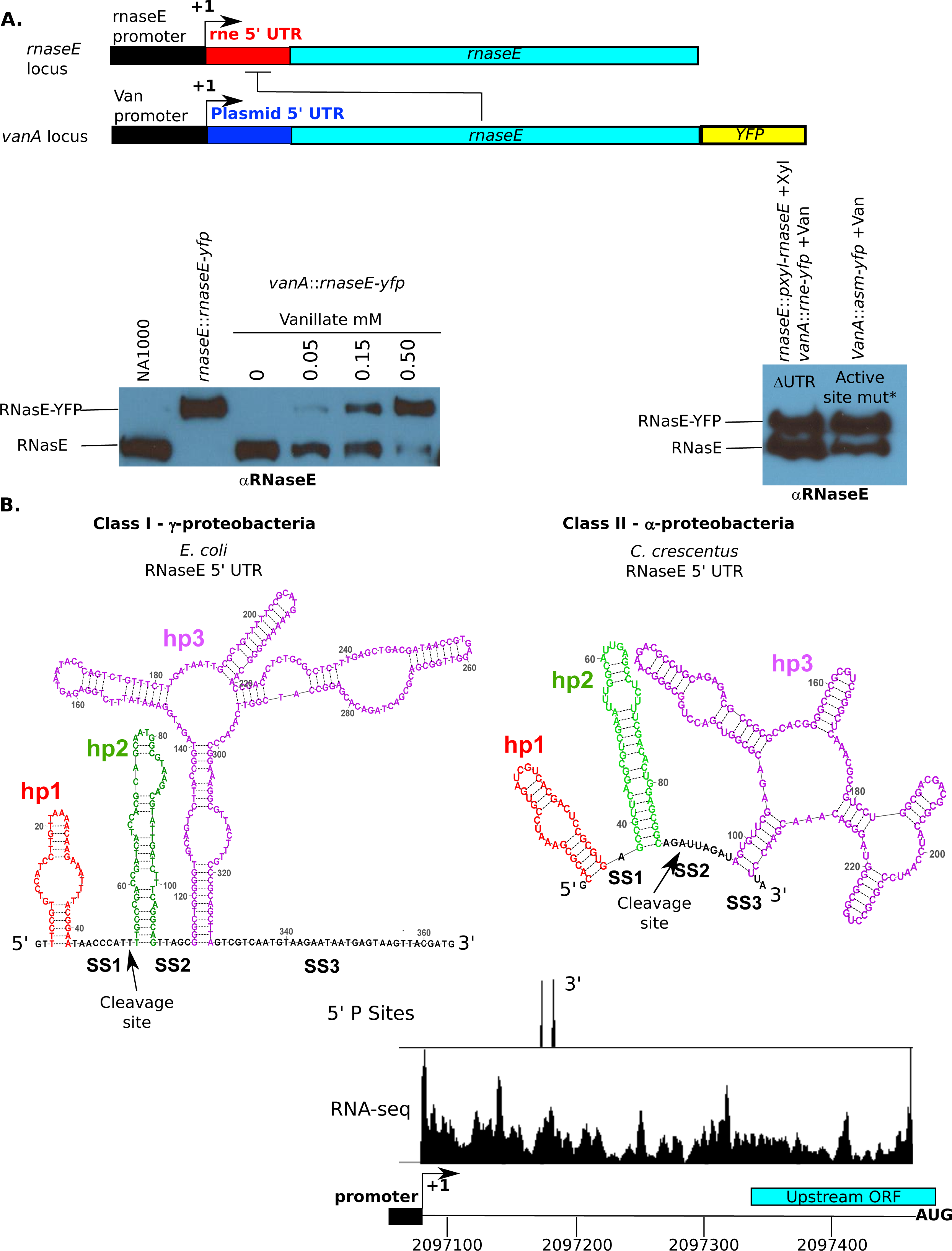
RNase E autoregulation requires activity on its 5’ UTR. A.) Western blot of RNase E in the indicated strains. Vanillate induced RNase E lacks the 5’ UTR. RNase E-YFP and RNase E bands are indicated. B.) Predicted mRNA secondary structures generated by turbofold for *C. crescentus* (left) and for *Escherichia coli* (right). RNA cleavage sites are mapped with arrows. RNA decay sites identification in the 5’ UTR. 5’ P-sites from(19) and RNA-seq data indicating the mRNA 5’ UTR from(42).

We investigated whether the *C. crescentus* RNase E had an annotated 5’ UTR structure in Rfam (12) but no 5’ UTR annotations existed outside of gamma-proteobacteria. To explore the 5’ UTR’s potential for structure formation, we extracted the 5’ UTR sequence from our RNA-seq based annotation (17) and performed secondary structure prediction using turbofold (18). This yielded a putative secondary structure that was similar to the *E. coli* 5’ UTR (Fig 3B), and that could be aligned to other α- and γ-proteobacterial RNase E 5’ UTRs (Fig S2), suggesting that the *C. crescentus* RNase E 5’ UTR and those in α-proteobacteria are likely a structural variant of the *E. coli* 5’ UTR. Secondary structure comparison between the γ- and α-proteobacteria show that the RNase E 5’ UTR likely exists with two different classes: Class I occurs in the γ-proteobacteria with a larger single stranded region I located between hairpins 1 and 2, while Class II occurs in the α-proteobacteria with a shorter single stranded region I and a larger single stranded region 2 (Fig S3 A). To identify *C. crescentus* 5’ RNA cleavage sites, we reanalyzed TAP dependent global 5’ end sequencing data which was used to identify 5’ PPP-containing transcription start sites to identify enriched 5’ P-cleavage sites (19) whereby we identified two RNA cleavage sites within the 5’ UTR, located in the single stranded region II (Fig 3B). The location of these cleavage sites differs from *E. coli*, where single stranded region I is the location of an RNase E cleavage site (Fig 3B). This suggests that the variation in the single stranded region may alter the preference where RNA cleavage by RNase E occurs between class I and II 5’ UTRs.

To test whether autoregulation impacts the cellular fitness we compared RNase E replacement strains expressing either RNase E-YFP with the natural 5’ UTR or a plasmid derived 5’ UTR (Fig 4A). In RNase E replacement strains the cells contain two copies of RNase E: the UTR variants were expressed from the *vanA* locus of the chromosome, while a xylose inducible promoter integration plasmid was introduced at the native *rne* locus, thereby the 5’ UTR variants are the only expressed copies of RNase E when grown in PYE (13). The strain harboring the 5’ UTR replaced with a plasmid 5’ UTR (JS38) led to a ∼2-fold higher RNase E-YFP levels than the strain harboring RNase E’s own 5’ UTR (JS249) (Fig 4A). When grown in PYE-vanillate conditions, we found that JS38 (106 minutes doubling time) had a decrease in growth rate compared to JS249 (94 minutes doubling time) (Fig 4A), suggesting that RNase E autoregulation promotes faster growth. While the magnitude in growth rate difference was small, we sought to assay fitness between the strains more sensitively by performing a growth competition experiment. In this competition experiment, a 50:50 ratio of each strain was incubated together and grown for multiple generations and the fraction of JS38 and JS249 cells in the population were measured. To measure the fraction of each strain in the population, we imaged the cells during mid-log phase of growth and measured their YFP intensities, which yielded two peaks in fluorescence intensity that could be fit with a dual gaussian curve-fit, and the area under each peak representing JS249 maxima (low YFP centered around 200 AU) and JS38 (high YFP centered around 350 AU) were used to calculate the ratio of each strain in the mixed culture (Fig 4B,C). Each day after incubating the mixed culture we observed a growing fraction of JS249 cells and a corresponding reduction in JS38 cells (Fig 4B). After three days of growth competition, we observed ∼12x more JS249 cells than JS38 cells, suggesting that autoregulation can promote a significant fitness advantage to *C. crescentus* cells (Fig 4 B, C).

**Figure 4.**
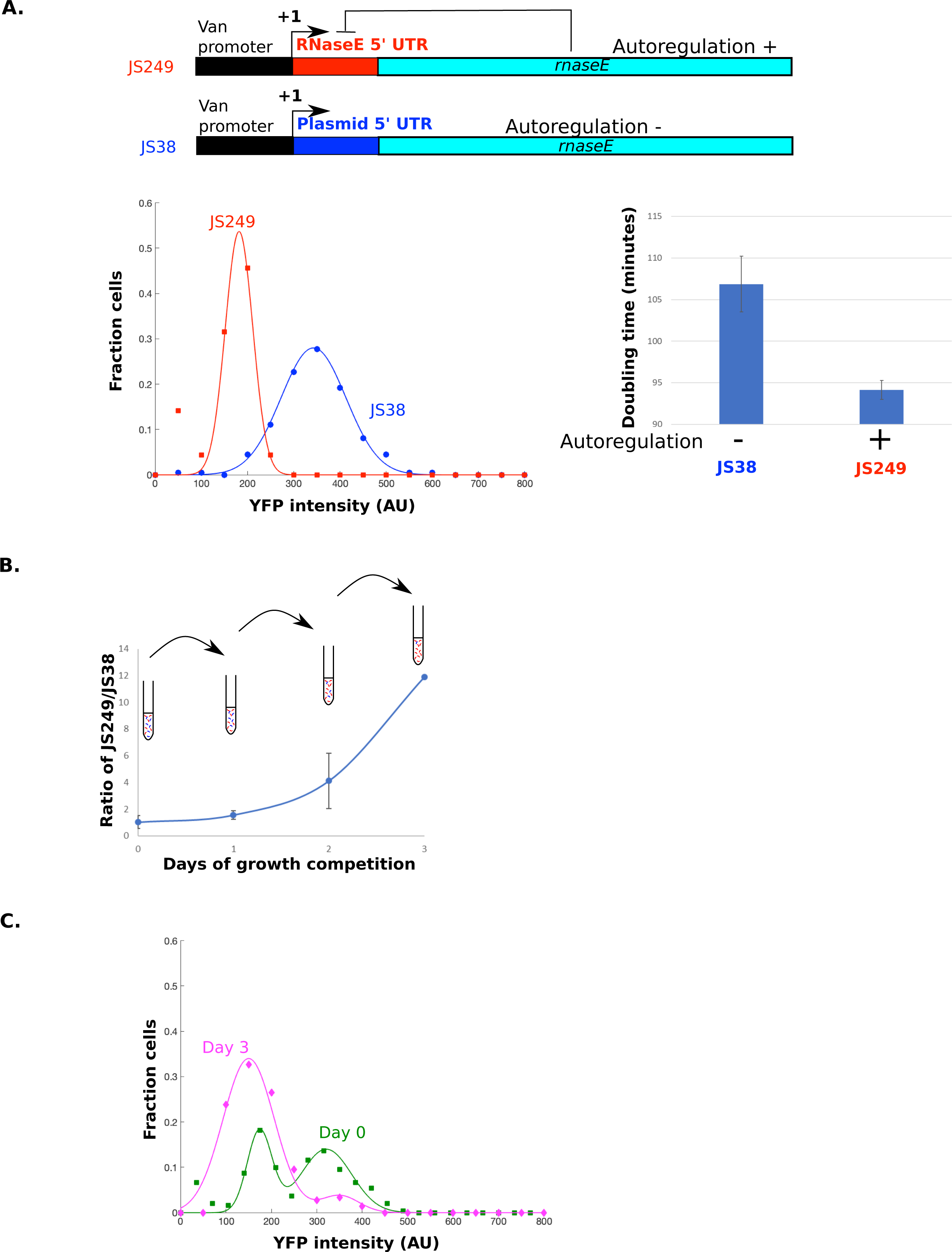
RNase E autoregulation contributes to cellular fitness. A.) Cartoon of strains JS249 containing the 5’ UTR, and JS38 which has a 5’ UTR from a plasmid expression system. YFP intensity distributions of strains JS249 and JS38, gaussian fits were performed with kaleidagraph. B.) Doubling times of JS38 and JS249 (left) as grown in PYE media. Growth competition of equally mixed cultures containing JS249 and JS38 after multiple days of growth (right). Error bars in both panels from 3 biological replicates. C.) YFP distributions of the mixed cultures at the beginning of the growth competition or after 3 days of culture. Dual gaussian fits were performed using kaleidagraph.

### RNase E’s intrinsically disordered region promotes phase separation and 5’ UTR cleavage

In *E. coli* and *C. crescentus*, the intrinsically disordered CTD (or IDR henceforth) of RNase E was found to be necessary for autoregulation (13, 14). While the IDR is also the scaffolding site for the RNA degradosome complex, we found an IDR mutant lacking the 3 degradosome proteins binding sites (Δ Aconitase, Δ RNase D, Δ PNPase) was capable of autoregulation (13), suggesting autoregulation does not require the formation of the RNA degradosome. We further dissected the N-terminal domain by dissecting it into its two sub-domains, the N-terminal S1 domain (amino acid 1-176) and the E/G catalytic core (amino acid 177-450) (13). We also observed that deletion of the N-terminal S1 domain, which binds RNA, or deletion of the catalytic E/G domain also abolished autoregulation (Fig S3C), suggesting a fully functional N-terminal domain is required for autoregulation. The IDR was found to be necessary and sufficient to form BR-bodies(13, 20), which allows RNase E to phase-separate with RNA and stimulates RNA decay activity *in vivo*(7, 13). To test whether the IDR promotes 5’ UTR cleavage *in vitro*, we purified full length RNase E and RNase E lacking the IDR. To confirm purified RNase Es were active we performed 9S ribosomal RNA processing assays with RNase E and RNase EΔIDR since RNase E is required for processing 9S rRNA to precursor 5S ribosomal RNA (p5S rRNA) (21). We observed that both RNase E and RNase EΔIDR processes 9S rRNA to p5S rRNA (Fig S3A), suggesting both proteins were functionally active. Next, we *in vitro* transcribed the rne 5’ UTR and tested its cleavage by RNase E. Under conditions with low enzyme to substrate ratio, initial RNA cleavage fragment size was observed to be consistent with the *in vivo* 5’P cleavage site [Fig 3B, S3B], which appears to be followed by subsequent cleavage events upon prolonged incubation.

To test the role of the IDR and impacts of phase-separation on RNase E autoregulation *in vitro*, we sought to incubate RNase E and RNase EΔIDR under conditions in which they were incubated above or below the critical concentration of phase separation. In prior work, *C. crescentus* RNase E’s IDR was incubated between 2-48 µM with poly-A RNA under a range of different monovalent salt concentrations, and it was observed to undergo phase-separation at a critical concentration >2 µM (Fig 5A) (13). Therefore, we incubated full length RNase E reactions at two concentrations, one concentration above the critical threshold for RNase E phase separation (6 µM RNase E), and one below the critical threshold for phase separation (1 µM RNase E) (Fig 5A). We confirmed these results with full length RNase E, where we observed that when incubated at 6 µM with its 5’ UTR RNA, it formed condensates under the microscope (Fig 5A); however, only a diffuse solution of RNase EΔIDR at 6 µM was observed in the presence of 5’ UTR RNA, in line with the role of the C-terminal IDR driving phase separation *in vivo* (13). Additionally, incubating 1 µM RNase E or RNase EΔIDR with 5’ UTR RNA did not lead to condensate formation (Fig 5A). We then performed single-turnover *in vitro* 5’ UTR RNA cleavage assays at both phase separating concentrations (6 µM) and non-phase separating conditions (1 µM) while keeping the enzyme/RNA ratio constant. We observed that at 1 µM RNase E cleaved the RNA slightly faster than RNase EΔIDR, as we observed increased amounts of RNA cleavage intermediates (Fig 5B), however, even after 30 minutes of incubation with RNase E the amount of full length 5’ UTR RNA remained high, and only a small fraction of cleavage products was detected (Fig S3B). When RNase E was incubated at 6 µM, its 5’ UTR was degraded rapidly, with only a small fraction of full-length RNA remaining after 15 minutes of incubation and the rest of the RNA being cleaved into decay fragments. In contrast, the RNase EΔIDR mutant incubated at 6 µM cleaved the RNA more slowly than full length, with the major population of RNA being uncleaved after 15 minutes of incubation, and a much lower fraction of RNA decay fragments accumulated than the full length. Taken together, this suggests that RNase E’s IDR accelerates 5’ UTR cleavage under conditions that promote condensation.

**Figure 5.**
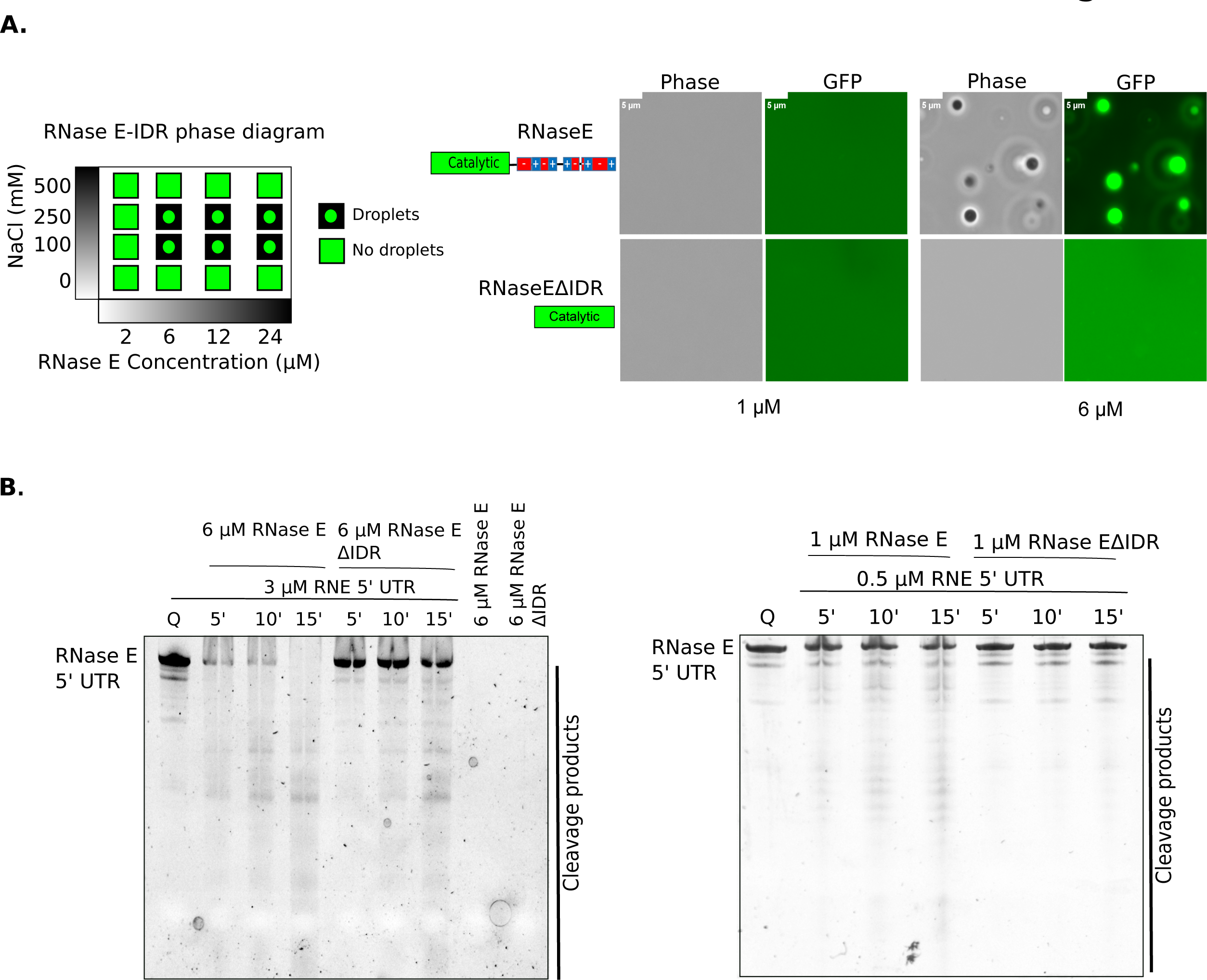
RNase E 5’ UTR cleavage is stimulated by the IDR. A.) Purified solutions of full-length RNase E containing the IDR (left) and the ΔIDR variant (right) after incubated with RNA for 30 minutes. Scale bar is 5 µm. B.) RNase E 5’ UTR cleavage assay. RNase E’s 5’ UTR was incubated with either full length RNase E or the ΔIDR variant for the indicated time periods before running on a 7% denaturing PAGE and stained by SYBR gold.

## Discussion

### RNase E autoregulation tunes RNase E levels for optimal growth

RNase E is typically the rate limiting enzyme controlling mRNA decay, so its protein levels must be carefully controlled to allow for the required mRNA decay activity for the cell. Negative autoregulation of RNase E appears to help fulfill this requirement by fine-tuning the amount of RNase E activity in the cell via cleavage of its own 5’ UTR (Figs 3,4,5). Levels of RNase E that are too low or too high fail to support cell growth (Figs 1,2), while mild overexpression of RNase E in cells lacking its native 5’ UTR leads to a more subtle but significant defect in cell growth and fitness (Fig 4). This suggests that even minor alterations of RNase E levels can negatively alter cell physiology and fitness. Therefore, RNase E’s 5’ UTR appears to be more broadly conserved outside the γ-proteobacteria to precisely control RNase E levels that fine-tunes its cellular concentrations and sets the cellular mRNA decay rate.

### RNase E condensates accelerate 5’ UTR cleavage

Past studies found that autoregulation *in vivo* required the 5’ UTR and the intrinsically disordered C-terminal domain of RNase E (14). This is the same IDR region of RNase E that is necessary and sufficient to phase-separate into BR-bodies, which are biomolecular condensates that promote faster RNA cleavage *in vivo* (7, 13). Interestingly, using *in vitro* reconstituted RNase E and RNase EΔIDR incubated above and below the critical concentration for phase-separation, we showed that the full-length version of RNase E can phase-separate with the 5’ UTR *in vitro* yielding faster 5’ UTR cleavage than RNase EΔIDR, while below the critical concentration the proteins were more similar in their 5’ UTR cleavage rates. This suggests that RNase E’s phase-separation with its 5’ UTR promotes more rapid RNA cleavage, perhaps by increasing the local concentration of RNA and RNase E within the condensate and may also explain why RNase E IDR deletion mutants have been found to globally slow mRNA decay (13, 22). We also observed previously that RNase E condensates are observable within seconds after the addition of RNA, suggesting that phase separation of BR-bodies is also a rapid process (20). Importantly, the RNA degradosome binding partner PNPase, which is a 3’ to 5’ exoribonuclease, was also shown to have increased exonucleolytic activity in the presence of RNase E droplets (23). This suggests that BR-bodies help coordinate the multi-step RNA decay process by concentrating essential mRNA decay enzymes with their mRNA substrates, not only to accelerate the rate-limiting step of RNA decay by RNase E, but also to promote the subsequent 3’ to 5’ exonucleolytic steps that complete mRNA decay, thereby preventing the pre-mature release of RNA decay intermediates. While *C. crescentus* provides an ideal model for the biochemical characterization of BR-bodies, the RNA degradosome machinery in multiple other bacteria and in mitochondria has been found to form foci *in vivo* (19, 24, 24–28), and which contain large IDRs, suggesting BR-bodies are likely widespread condensates used for mRNA decay. Biomolecular condensation therefore provides a general organizational strategy for bacteria to organize their biochemical pathways in the absence of membrane-bound organelles. Indeed, many bacterial enzymes from various multi-step biochemical pathways have been found to phase-separate and form biomolecular condensates (20, 28–41) suggesting that this strategy of subcellular organization may be utilized to organize a variety of other biochemical pathways in bacteria.

## Materials and Methods

### Cell growth

All strains used in this study were derived from the wild-type strain NA1000, and were grown at 28°C in peptone-yeast extract (PYE) medium or M2 minimal medium supplemented with 0.2% D-glucose (M2G). When appropriate, the indicated concentration of Vanillate (5 µM), Xylose (0.2%), gentamycin (0.5 μg/mL), kanamycin (5 μg/mL), chloramphenicol (2 μg/mL), oxytetracycline (2µg/ml), spectinomycin (25 μg/mL), and/or streptomycin (5 μg/mL) were added. Strains were analyzed at mid-exponential phase of growth (OD 0.3-0.6). Optical density was measured at 600 nm in a cuvette using a Nanodrop 2000C spectrophotometer. For depletion, a strain containing a xylose inducible copy of RNase E was first grown in media containing xylose overnight, then washed 3 times with 1mL growth media, and resuspended in growth media without xylose and grown for 8 hours - overnight. Log-phase cultures were then used for any downstream application. For overexpression, overexpression strains containing a xylose inducible copy of RNase E variant were first grown in media without xylose overnight, then induced with xylose for 3.5-4 hours. Autoregulation test strains containing a copy of *rne-yfp* integrated at the *vanA* locus were grown overnight in PYE, then induced with the indicated vanillate concentration for six hours. Replacements strains containing a xylose inducible copy of RNase E and a Vanillate inducible test construct were first grown in media containing xylose overnight. Log-phase cells were washed 3 times with growth plain media and used to inoculate resuspended in growth media containing Vanillate, diluted, and grown overnight. Log-phase cultures were then used for any downstream application.

### Serial dilutions assay

For serial dilution assays, cells were grown overnight in PYE-kan media supplied with 0.2% xylose. Log-phase cells were washed 3 times with PYE media and diluted in PYE without xylose to an OD 0.05. Serial dilutions were then performed and 5 µls of the selected dilutions were spotted on PYE-kan plates with and without xylose and incubated at 28°C. For overexpression strains, the cells were grown in PYE-kan media without xylose overnight and log-phase cells were diluted in PYE-kan without xylose to an OD 0.05. Serial dilutions were then performed and 5 µls of the selected dilutions were spotted on PYE-kan plates with and without xylose and incubated at 28°C. For replacements strains, cells were first grown in PYE-kan-gent media supplied with 0.2% xylose overnight. Log-phase cells were washed 3 times with plain PYE media and resuspended in plain PYE media to an OD 0.05. Serial dilutions were then performed and 5 µls of the selected dilutions were spotted on PYE-kan-Gent, PYE-kan-Gent-xylose and PYE-kan-Gent-vanillate plates and incubated at 28°C.

### Western blots

For determining the optimal RNE depletion time, JS8 cells were grown in PYE-kan media containing 0.2% xylose overnight, then washed 3 times with PYE media. The washed cells were used to inoculate 30 ml of PYE media without xylose. Samples were taken at 1,2,3,4,8 hours after xylose removal. The cells were pelleted and resuspended in 250μl of 1x laemmli buffer for each 1.0 OD_600_ unit. For determining the optimal RNE overexpression time, JS89 cells were grown in PYE media overnight, log-phase cells were used to inoculate 30 ml of PYE-kan media with 0.2% xylose. Samples were taken at 1,2,3,4 hours after xylose addition. The cells were pelleted and resuspended in 250μl of 1x laemmli buffer for each 1.0 OD_600_ unit. The western blotting was performed as in(13) using (1:1000) dilution of α-RNE antibodies.

### 5’ UTR structure prediction

The 5’ UTR of RNase E was extracted from(17), and the sequence from the +1 site to the start codon was placed into turbofold(18) for secondary structure prediction. As a control, the *E. coli* 5’ UTR of RNase E from (11) was predicted in the same manner.

### 5’ P site identification

For determining 5’ P sites in the 5’ UTR, we analyzed the TAP-samples from(19) that occurred within the 5’ UTR region.

### Growth competition experiments

The strains used in the experiments included the strain with a mutated 5’ UTR (JS38) and one with a WT 5’ UTR (JS249). JS38 has the functional RNase E gene under the xylose-inducible promoter and a YFP-tagged RNase E with a mutated 5’ UTR under the vanillate-inducible promoter. JS249 also has a functional RNase E gene under the xylose-inducible promoter, but it has a YFP-tagged RNase E with the wild-type 5’ UTR region under the vanillate-inducible promoter.

A competition experiment was conducted to compare the growth efficiencies of the two strains. To start, both strains were grown in five-milliliter overnight cultures at 28°C in PYE with xylose and antibiotics Kanamycin (Kan) and Gentamycin (Gent). The next day, cells were pelleted and washed to remove the xylose and then resuspended in one milliliter of PYE. An optical density (OD_600_) was taken for each culture, and the cultures were diluted to 0.05 in five-milliliter cultures of PYE/Kan/Gent and induced for 6 hours with vanillate in 28°C shaker. After cells grew for 6 hours, optical OD_600_ was taken for each culture, and the culture with a greater OD_600_ was diluted to match the culture with the lower initial OD_600_. From here, a mixed culture of JS38 and JS249 was created. Cells were spotted on agarose pads and imaged through fluorescence microscopy with a YFP filter cube and exposure time of 700ms for both the cultures. These images represent the Day 0 images. Serial dilutions of JS38, JS249, and JS38 vs. JS249 were made and grown overnight with vanillate to ensure log phase imaging for the following day. Images were taken until Day 3 for Trial 1 and Day 2 for Trial 2. A master mix of PYE/Kan/Gent/Van was made to be used for cultures starting with the 6-hour induction on Day 0 and the controls were imaged every day.

### Competition strain fluorescence measurements

The experiment resulted in six images of each strain (JS38, JS249, and JS38 vs. JS249) from each day. The fluorescence intensities were analyzed using the program ImageJ. The background intensity of each image, found using MicrobeJ, was subtracted from the fluorescent intensities. The resulting fluorescent intensities were used for the analysis.

Due to overlap between the median fluorescent intensities of the two strains, cells from the mixed strains could not be clearly separated by fluorescence intensities. Instead, using the program, KaleidaGraph, a Gaussian curve fit was created for both JS38 and JS249, and the functions of these graphs were summed to create a double-Gaussian curve fit for the mixes each day. In each of the graphs, the values were normalized to account for differences in the number of cells imaged per day. The ratio of the areas under the curve was calculated, JS249:JS38 by taking the integral of both parts of the double-Gaussian functions separately from zero to one thousand. The expected result is a 1:1 ratio on Day 0 and an increase in the ratio each following day.

### Competition strain growth rate measurements

The experiment was conducted in a triplicate to ensure reproducibility. JS38 and JS249 were grown in five-milliliter overnight cultures containing PYE/Kan/Gent and xylose at 28°C. The cells were washed to remove the xylose and induced overnight in serial dilutions with Van. The next day, an OD_600_ was measured, and the cells in log phase were used to inoculate 6 50-milliliter cultures, creating an OD_600_ of around 0.05. OD_600_ time points were taken for each of the 6 flasks every 90 minutes, and the data was used to analyze the doubling time of each strain. A master mix of PYE/Kan/Gent/Van was used starting with the overnight in Van and used for the cultures, as well as the media for blanking while taking OD_600_.

### RNase E purification

Full length RNase E gene was amplified from *C. crescentus* genome using VN106F and VN107R primers and cloned into pET-GFP vector through ligation independent cloning (LIC). The sequence verified plasmid was transformed into BL21(DE3) cells and the resultant colonies were inoculated into LB media (50 mL) with 30 µg/mL Kanamycin and grown overnight at 37°C, 200 rpm. The saturated culture was reinoculated into 1.5-liter LB media containing 30 µg/mL Kanamycin and grown at 37°C, 200 rpm until the OD reaches ∼0.6. RNase E expression was induced with 0.5 mM IPTG at 37°C, 140 rpm for 3 hours. The cells were harvested at 5000 rpm for 15 mins and resuspended in 20mL of Lysis buffer (20mM Tris pH 7.4, 500mM NaCl, 10% glycerol, 10mM imidazole, 10 µg/mL DNase I). The cells were lysed in Sonicator at 4°C with 5 sec on time and 10 sec off time for 3 minutes. The lysate was centrifuged at 4°C, 14000 rpm for 45 minutes and the resultant supernatant was passed through pre-equilibrated Ni-NTA resin for binding of GFP-RNase E-His to the resin. After washing the protein bound resin with 5 column volumes each of low salt buffer (20mM Tris pH 7.4, 150mM NaCl, 5% glycerol, 10mM imidazole) and high salt buffer (20mM Tris pH 7.4, 1000mM NaCl, 5% glycerol, 10mM imidazole), the protein was eluted in elution buffer (20mM Tris pH 7.4, 150mM NaCl, 5% glycerol, 250mM imidazole). The protein was concentrated using amicon concentrator with 30 kDa cutoff and the concentrated protein was passed through S200 Sephadex size exclusion column in SEC buffer (20mM Tris pH 7.4, 250mM NaCl, 2% glycerol) for further purification. The resultant pure protein was concentrated to 15 mg/mL and stored in −80°C.

The GFP RNase E Δ IDR (RNase E Δ IDR) was the proteolyzed fragment recovered from size exclusion chromatography while performing GFP RNase E full length (RNase E) purification.

### *In vitro* transcription of RNase E 5’ UTR and 9S rRNA

The plasmid containing RNase E 5’ UTR (pVN053) was linearized using Nhe1 restriction enzyme, which served as template in *in vitro* transcription (IVT). The IVT reaction mixture contained 2.7mg of linearized plasmid, 21µg homemade T7 RNA polymerase, 2.5mM NTPs, 1X reaction buffer (50mM Tris-Cl pH 7.4, 15mM MgCl_2_, 5mM DTT, 2mM spermidine), 0.01 U/µL PPase in a total volume of 1000 µL and the reaction was carried out at 37°C for 4 hours. The transcribed RNA loaded onto 7% Urea gel was extracted using phenol-chloroform and ethanol precipitation methods. The precipitated RNA was resuspended in nuclease free water for carrying out RNase E cleavage assays. The template for 9S rRNA *in vitro* transcription was created by PCR amplification (using VN108 F and VN109 R) consisting of T7 promoter and 9S rRNA sequence.

### pCp Cy5 labeling of RNase E 5’ UTR

*In vitro* transcribed RNase E 5’ UTR (4.5 µM) was incubated with 33 µM of pCp Cy5 (Jena Bioscience #NU-1706-CY5) in a total reaction volume of 100 µL consisting of 50 units of T4 RNA ligase 1 (NEB #M0204S), 1mM ATP, 10% DMSO1, 1X T4 RNA ligase reaction buffer (NEB #B0216S) for 16 hours at 16°C. Following the reaction, T4 RNA ligase 1 was heat inactivated at 65°C for 15 minutes and the reaction mixture was subjected to Phenol-Chloroform extraction to remove the enzyme. The unincorporated pCp Cy5 was removed by passing the labeled RNA through Sephadex G-50 column (Cytiva # 28903408) and eluted in nuclease free water.

### *In vitro* RNase E condensate formation assay

6 µM of GFP RNase E or GFP RNase E Δ IDR were incubated with 25ng/µL RNase E 5’ UTR - Cy5 RNA in 20mM Tris pH 7.4, 100mM NaCl, 1mM DTT buffer in a total reaction volume of 10 µL for 30 minutes at room temperature. Entire 10 µL was spotted on a slide and covered with coverslip before imaging with Nikon Eclipse NI-E with CoolSNAP MYO-CCD camera and 100x Oil CFI Plan Fluor (Nikon) objective under phase contrast, GFP and Red channels with exposures of 30ms, 50ms and 50ms respectively.

### *In vitro* 9S rRNA processing assay

0.1 µM RNase E or RNase E Δ IDR was incubated with 1 µM of 9s rRNA in 20mM Tris pH 7.4, 150mM NaCl, 2% glycerol, 1mM DTT, 100 µM MgCl_2_ buffer in a total reaction volume of 40 µL at 28°C. At the end of each time point, 10 µL of the reaction volume was mixed with 15 µL of stop buffer (95% formamide, 50mM EDTA, 0.1% SDS, 0.025% bromophenol blue, 0.025% xylene cyanol). The samples were heated at 90°C for 3 mins and resolved on 7% Urea-acrylamide gel, stained with 1X SYBR Gold nucleic acid stain and scanned using GE Typhoon FLA 9000 gel scanner.

### *In vitro* RNase E 5’ UTR cleavage assay

1 µM of GFP RNase E or GFP RNase E Δ IDR was incubated with 0.5 µM of RNase E 5’ UTR in 1X reaction buffer (20 mM Tris pH 7.4, 150mM NaCl, 2% glycerol, 1mM DTT, 100 µM MgCl_2_) in a total reaction volume of 40 µL. At the end of each time point, 10 µL of this reaction mixture was added to 15 µL of stop buffer (95% formamide, 50mM EDTA, 0.1% SDS, 0.025% bromophenol blue, 0.025% xylene cyanol). The samples were heated at 90°C for 3 minutes and loaded onto 7% urea-acrylamide gel. The gel was stained with 1X SYBR Gold nucleic acid stain (Thermo Fisher #S11494) for 20 minutes and scanned using Thermofisher iBright imaging system.

For the cleavage assay under condensate forming condition, 6 µM RNase E or RNase E Δ IDR was incubated with 3 µM RNase E 5’ UTR and the reaction was carried out as mentioned above. At the end of each time point, the reaction is diluted 50 times so that the RNA loaded in each well is 100ng.

### Plasmid Construction

#### pVN053

RNase E 5’UTR along with the first 150 bases of coding sequence was amplified from *C. crescentus* genome using VN082F & VN083R primers. The amplified sequence was ligated into pMCS-2 vector under T7 promoter through Gibson assembly. Positive colonies were selected on LB-Kan plate followed by sanger sequencing.

#### pVN059

Full length RNase E gene was amplified from *C. crescentus* genome using VN106F and VN107R primers and cloned into pET-GFP vector through ligation independent cloning (LIC).

#### pBXMCS-2 RNE YFP

RNase E-YFP from the pVRNEYFPC-4 plasmid was digested with NdeI and NheI and ligated into pBXMCS-2 cut with NdeI and XbaI. The plasmid was sequence verified (genewiz) before electroporation into Caulobacter.

#### Strain Construction

##### JS8: NA1000 RNE::pXRNEssrAC KanR

The strain was generated as described before (13).

##### JS49: NA1000 vanA::FL-RNE-YFP GentR

The strain was generated as described before (13).

##### JS61: NA1000 vanA::RNE(ΔE/G)-YFP Gent^R^

The strain was generated as described before (13).

##### JS62: NA1000 vanA::RNE(DRhlBBS)-YFP Gent^R^

The strain was generated as described before (13).

##### JS89

The strain was generated by transforming pBXMCS-2 – RNE YFP plasmid into NA1000 cells and the positive colonies were selected on PYE-kan medium followed by YFP fluorescence in the cells.

##### JS249

To generate JS249, we inserted the 5’ UTR into the pVRNEYFPC-4 plasmid via Gibson assembly. The 5’ UTR was amplified from the NA1000 chromosome using RNE5’UTR_F and RNE5’UTR_R primers and the pVRNEYFPC-4plasmid containing RNaseE-YFP was amplified using PVRNEYFPC-4_F and PVRNEYFPC-4_R primers. We then performed Gibson assembly using both fragments, transformed into DH10B cells, and selected on LB-Kan plates. Colonies were then screened for the RNase E 5’ UTR and sequence verified. The resulting pVRNE5’UTR-RNEYFPC-4 plasmid was then transformed into NA1000 cells by electroporation and selected on PYE-gent plates. GentR colonies were then subjected to phage transduction using JS8 lysates and selected on PYE-gent-kan-xylose plates. The resulting colonies were phenotyped for xylose and vanillate dependent growth.

##### JS 293: NA1000 vanA::RNE(ASMmut2)YFPC Gent^R^

The strain was generated as described before (13).

##### Table of oligos

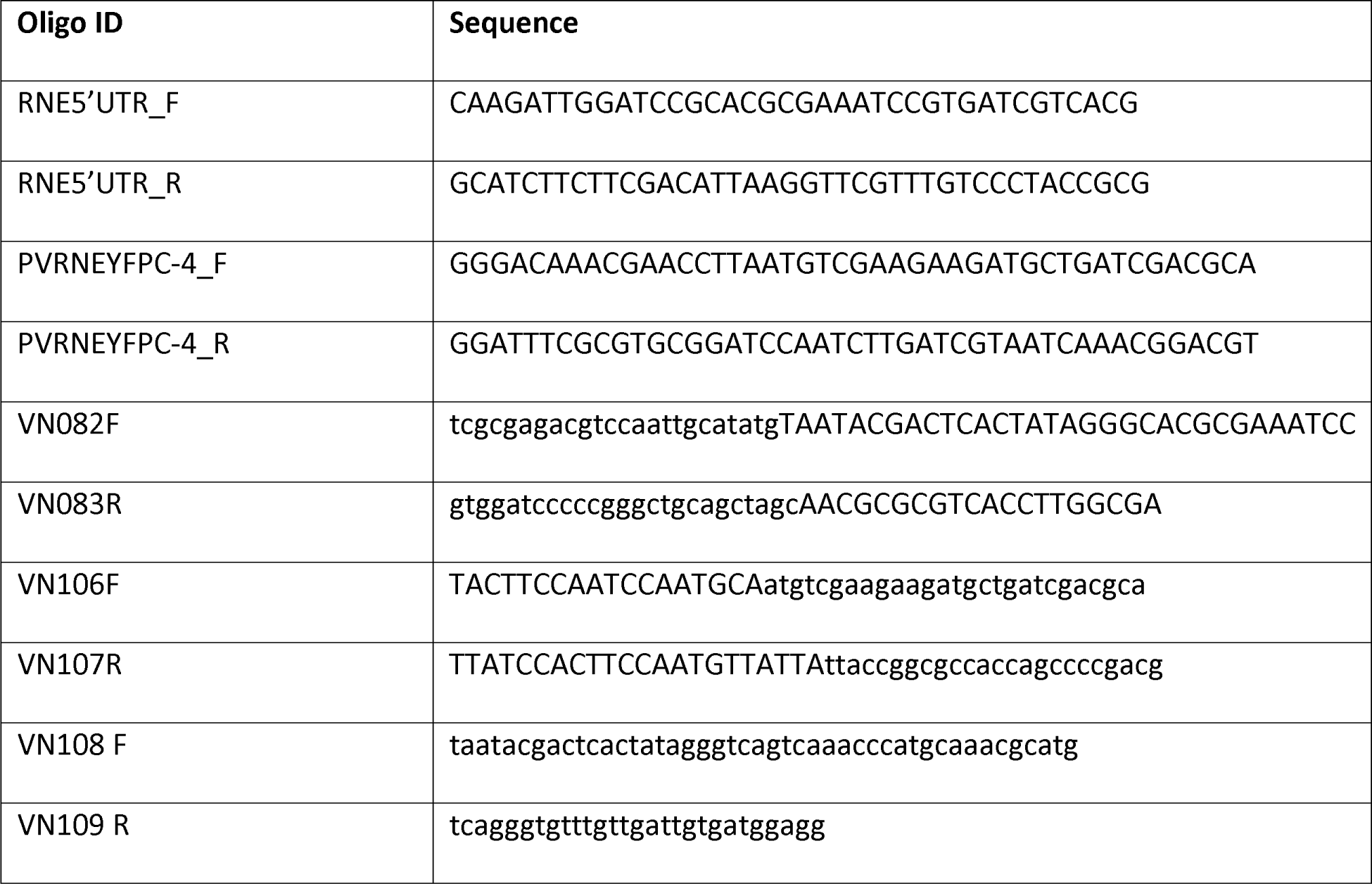

##### Table of Strains used

| Strain name | Genotype |
| --- | --- |
| JS8 | NA1000 RNE::pXRNEssrAC Kan<sup>R</sup> |
| JS49 | NA1000 vanA::FL-RNE-YFP Gent<sup>R</sup> |
| JS61 | NA1000 vanA::RNE(ΔE/G)-YFP Gent<sup>R</sup> |
| JS62 | NA1000 vanA::RNEΔS1-YFP Gent<sup>R</sup> |
| JS89 | NA1000+pBXMCS2 with FLRNE-YFP Kan<sup>R</sup> |
| JS221 | NA1000 vanA::RNase E ΔIDR-YFP integrated |
|  | +RNE::pXRNEssrAC-2 Gent<sup>R</sup> Kan<sup>R</sup> |
| JS249 | NA1000 vanA::RNE 5'UTR-YFP +<br>RNE::pXRNEssrAC-2 Gent<sup>R</sup> Kan<sup>R</sup> |
| JS 293 | NA1000 vanA::RNE(ASMmut2)YFPC Gent<sup>R</sup> |

## Author Contributions

JMS conceived the project. VN performed protein purifications, *in vitro* transcriptions, RNA labeling, RNA cleavage experiments and phase separation assays. NA and AV performed cell biology experiments and functional experiments. AV performed the growth competition experiments. KD performed *in vitro* transcription. JMS and VN wrote the paper.

## Acknowledgements

The authors thank Wayne State University startup funds to JMS. Research reported in this publication was supported by NIGMS of the National Institutes of Health under award number R35GM124733.

## References

1. Carpousis AJ, Luisi BF, McDowall KJ. 2009. Chapter 3 Endonucleolytic Initiation of mRNA Decay in Escherichia coli, p. 91–135. In Progress in Molecular Biology and Translational Science. Academic Press.

2. Mackie GA. 2013. RNase E: at the interface of bacterial RNA processing and decay. 1. Nat Rev Microbiol 11:45–57.

3. Chao Y, Li L, Girodat D, Förstner KU, Said N, Corcoran C, Śmiga M, Papenfort K, Reinhardt R, Wieden H-J, Luisi BF, Vogel J. 2017. In Vivo Cleavage Map Illuminates the Central Role of RNase E in Coding and Non-coding RNA Pathways. Molecular Cell 65:39–51.

4. Deutscher MP. 2006. Degradation of RNA in bacteria: comparison of mRNA and stable RNA. Nucleic Acids Research 34:659–666.

5. Kushner SR. 2002. mRNA Decay in Escherichia coli Comes of Age. Journal of Bacteriology 184:4658–4665.

6. Ono M, Kuwano M. 1979. A conditional lethal mutation in an Escherichia coli strain with a longer chemical lifetime of messenger RNA. Journal of Molecular Biology 129:343–357.

7. Al-Husini N, Tomares DT, Pfaffenberger ZJ, Muthunayake NS, Samad MA, Zuo T, Bitar O, Aretakis JR, Bharmal M-HM, Gega A, Biteen JS, Childers WS, Schrader JM. 2020. BR-Bodies Provide Selectively Permeable Condensates that Stimulate mRNA Decay and Prevent Release of Decay Intermediates. Molecular Cell 78:670–682.e8.

8. Schuck A, Diwa A, Belasco JG. 2009. RNase E autoregulates its synthesis in Escherichia coli by binding directly to a stem-loop in the rne 5′ untranslated region. Molecular Microbiology 72:470–478.

9. Jain C, Belasco JG. 1995. RNase E autoregulates its synthesis by controlling the degradation rate of its own mRNA in Escherichia coli: unusual sensitivity of the rne transcript to RNase E activity. Genes Dev 9:84–96.

10. Sousa S, Marchand I, Dreyfus M. 2001. Autoregulation allows Escherichia coli RNase E to adjust continuously its synthesis to that of its substrates. Molecular Microbiology 42:867–878.

11. Diwa A, Bricker AL, Jain C, Belasco JG. 2000. An evolutionarily conserved RNA stem–loop functions as a sensor that directs feedback regulation of RNase E gene expression. Genes Dev 14:1249–1260.

12. Kalvari I, Nawrocki EP, Ontiveros-Palacios N, Argasinska J, Lamkiewicz K, Marz M, Griffiths-Jones S, Toffano-Nioche C, Gautheret D, Weinberg Z, Rivas E, Eddy SR, Finn RD, Bateman A, Petrov AI. 2021. Rfam 14: expanded coverage of metagenomic, viral and microRNA families. Nucleic Acids Research 49:D192–D200.

13. Al-Husini N, Tomares DT, Bitar O, Childers WS, Schrader JM. 2018. α-Proteobacterial RNA Degradosomes Assemble Liquid-Liquid Phase-Separated RNP Bodies. Molecular Cell 71:1027–1039.e14.

14. Jiang X, Diwa A, Belasco JG. 2000. Regions of RNase E Important for 5′-End-Dependent RNA Cleavage and Autoregulated Synthesis. Journal of Bacteriology 182:2468–2475.

15. Christen B, Abeliuk E, Collier JM, Kalogeraki VS, Passarelli B, Coller JA, Fero MJ, McAdams HH, Shapiro L. 2011. The essential genome of a bacterium. Molecular Systems Biology 7:528.

16. Mudd EA, Higgins CF. 1993. Escherichia coli endoribonuclease RNase E: autoregulation of expression and site-specific cleavage of mRNA. Molecular Microbiology 9:557–568.

17. Bharmal M-H, Aretakis JR, Schrader JM. 2020. An Improved Caulobacter crescentus Operon Annotation Based on Transcriptome Data. Microbiology Resource Announcements 9:10.1128/mra.01025-20.

18. Tan Z, Fu Y, Sharma G, Mathews DH. 2017. TurboFold II: RNA structural alignment and secondary structure prediction informed by multiple homologs. Nucleic Acids Research 45:11570–11581.

19. Zhou B, Schrader JM, Kalogeraki VS, Abeliuk E, Dinh CB, Pham JQ, Cui ZZ, Dill DL, McAdams HH, Shapiro L. 2015. The Global Regulatory Architecture of Transcription during the Caulobacter Cell Cycle. PLOS Genetics 11:e1004831.

20. Nandana V, Rathnayaka-Mudiyanselage IW, Muthunayake NS, Hatami A, Mousseau CB, Ortiz-Rodríguez LA, Vaishnav J, Collins M, Gega A, Mallikaarachchi KS, Yassine H, Ghosh A, Biteen JS, Zhu Y, Champion MM, Childers WS, Schrader JM. 2023. The BR-body proteome contains a complex network of protein-protein and protein-RNA interactions. Cell Reports 42.

21. Hardwick SW, Chan VSY, Broadhurst RW, Luisi BF. 2011. An RNA degradosome assembly in Caulobacter crescentus. Nucleic Acids Res 39:1449–1459.

22. Lopez PJ, Marchand I, Joyce SA, Dreyfus M. 1999. The C-terminal half of RNase E, which organizes the Escherichia coli degradosome, participates in mRNA degradation but not rRNA processing in vivo. Molecular Microbiology 33:188–199.

23. Collins M, Tomares DT, Schrader JM, Childers WS. 2022. RNase E biomolecular condensates stimulate PNPase activity. bioRxiv 10.1101/2022.06.06.495039.

24. Griego A, Douché T, Gianetto QG, Matondo M, Manina G. 2022. RNase E and HupB dynamics foster mycobacterial cell homeostasis and fitness. iScience 25:104233.

25. Muthunayake NS, Tomares DT, Childers WS, Schrader JM. Phase-separated bacterial ribonucleoprotein bodies organize mRNA decay. WIREs RNA n/a:e1599.

26. Strahl H, Turlan C, Khalid S, Bond PJ, Kebalo J-M, Peyron P, Poljak L, Bouvier M, Hamoen L, Luisi BF, Carpousis AJ. 2015. Membrane Recognition and Dynamics of the RNA Degradosome. PLOS Genetics 11:e1004961.

27. Hamouche L, Billaudeau C, Rocca A, Chastanet A, Ngo S, Laalami S, Putzer H. 2020. Dynamic Membrane Localization of RNase Y in Bacillus subtilis. mBio 11.

28. Tejada-Arranz A, Galtier E, Mortaji LE, Turlin E, Ershov D, Reuse HD. 2020. The RNase J-Based RNA Degradosome Is Compartmentalized in the Gastric Pathogen Helicobacter pylori. mBio 11.

29. Nandana V, Schrader JM. 2021. Roles of liquid–liquid phase separation in bacterial RNA metabolism. Current Opinion in Microbiology 61:91–98.

30. Saurabh S, Chong TN, Bayas C, Dahlberg PD, Cartwright HN, Moerner WE, Shapiro L. 2022. ATP-responsive biomolecular condensates tune bacterial kinase signaling. Science Advances 8:eabm6570.

31. Lasker K, von Diezmann L, Zhou X, Ahrens DG, Mann TH, Moerner WE, Shapiro L. 2020. Selective sequestration of signalling proteins in a membraneless organelle reinforces the spatial regulation of asymmetry in Caulobacter crescentus. 3. Nature Microbiology 5:418–429.

32. Krypotou E, Townsend GE, Gao X, Tachiyama S, Liu J, Pokorzynski ND, Goodman AL, Groisman EA. 2023. Bacteria require phase separation for fitness in the mammalian gut. Science 379:1149–1156.

33. Goldberger O, Szoke T, Nussbaum-Shochat A, Amster-Choder O. 2022. Heterotypic phase separation of Hfq is linked to its roles as an RNA chaperone. Cell Reports 41:111881.

34. Ladouceur A-M, Parmar BS, Biedzinski S, Wall J, Tope SG, Cohn D, Kim A, Soubry N, Reyes-Lamothe R, Weber SC. 2020. Clusters of bacterial RNA polymerase are biomolecular condensates that assemble through liquid–liquid phase separation. PNAS 117:18540–18549.

35. Harami GM, Kovács ZJ, Pancsa R, Pálinkás J, Baráth V, Tárnok K, Málnási-Csizmadia A, Kovács M. 2020. Phase separation by ssDNA binding protein controlled via protein−protein and protein−DNA interactions. Proceedings of the National Academy of Sciences 117:26206–26217.

36. McQuail J, Switzer A, Burchell L, Wigneshweraraj S. 2020. The RNA-binding protein Hfq assembles into foci-like structures in nitrogen starved Escherichia coli. Journal of Biological Chemistry 295:12355–12367.

37. Azaldegui CA, Vecchiarelli AG, Biteen JS. 2021. The emergence of phase separation as an organizing principle in bacteria. Biophysical Journal 120:1123–1138.

38. Heinkel F, Abraham L, Ko M, Chao J, Bach H, Hui LT, Li H, Zhu M, Ling YM, Rogalski JC, Scurll J, Bui JM, Mayor T, Gold MR, Chou KC, Av-Gay Y, McIntosh LP, Gsponer J. 2019. Phase separation and clustering of an ABC transporter in Mycobacterium tuberculosis. Proceedings of the National Academy of Sciences 116:16326–16331.

39. Ramm B, Schumacher D, Harms A, Heermann T, Klos P, Müller F, Schwille P, Søgaard-Andersen L. 2023. Biomolecular condensate drives polymerization and bundling of the bacterial tubulin FtsZ to regulate cell division. 1. Nat Commun 14:3825.

40. Tan W, Cheng S, Li Y, Li X-Y, Lu N, Sun J, Tang G, Yang Y, Cai K, Li X, Ou X, Gao X, Zhao G-P, Childers WS, Zhao W. 2022. Phase separation modulates the assembly and dynamics of a polarity-related scaffold-signaling hub. 1. Nat Commun 13:7181.

41. Guilhas B, Walter J-C, Rech J, David G, Walliser NO, Palmeri J, Mathieu-Demaziere C, Parmeggiani A, Bouet J-Y, Le Gall A, Nollmann M. 2020. ATP-Driven Separation of Liquid Phase Condensates in Bacteria. Molecular Cell 79:293–303.e4.

42. Schrader JM, Zhou B, Li G-W, Lasker K, Childers WS, Williams B, Long T, Crosson S, McAdams HH, Weissman JS, Shapiro L. 2014. The Coding and Noncoding Architecture of the Caulobacter crescentus Genome. PLOS Genetics 10:e1004463.

43. Ducret A, Quardokus EM, Brun YV. 2016. MicrobeJ, a tool for high throughput bacterial cell detection and quantitative analysis. 7. Nat Microbiol 1:1–7.

