## Supplemental information for "*Caulobacter crescentus* RNase E condensation contributes to autoregulation and fitness"

### Figure S1

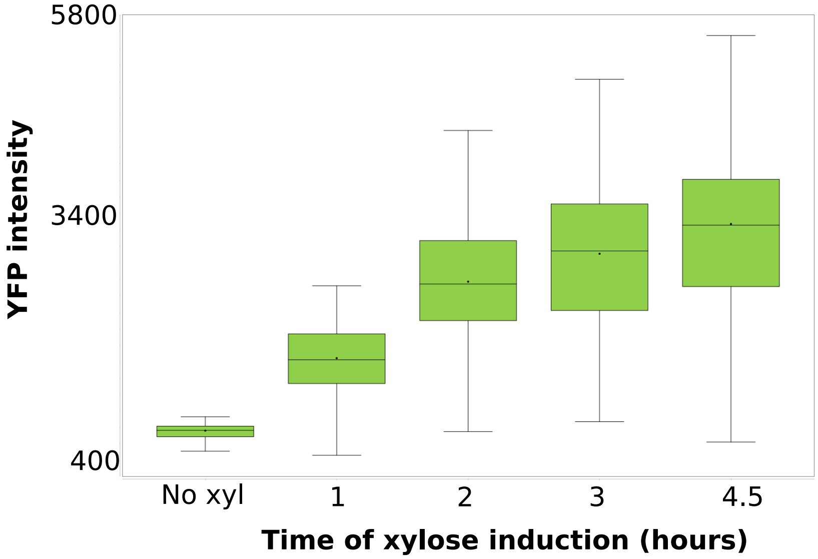
5800
3400
400
YFP intensity
No xyl
1
2
3
4.5
Time of xylose induction (hours)

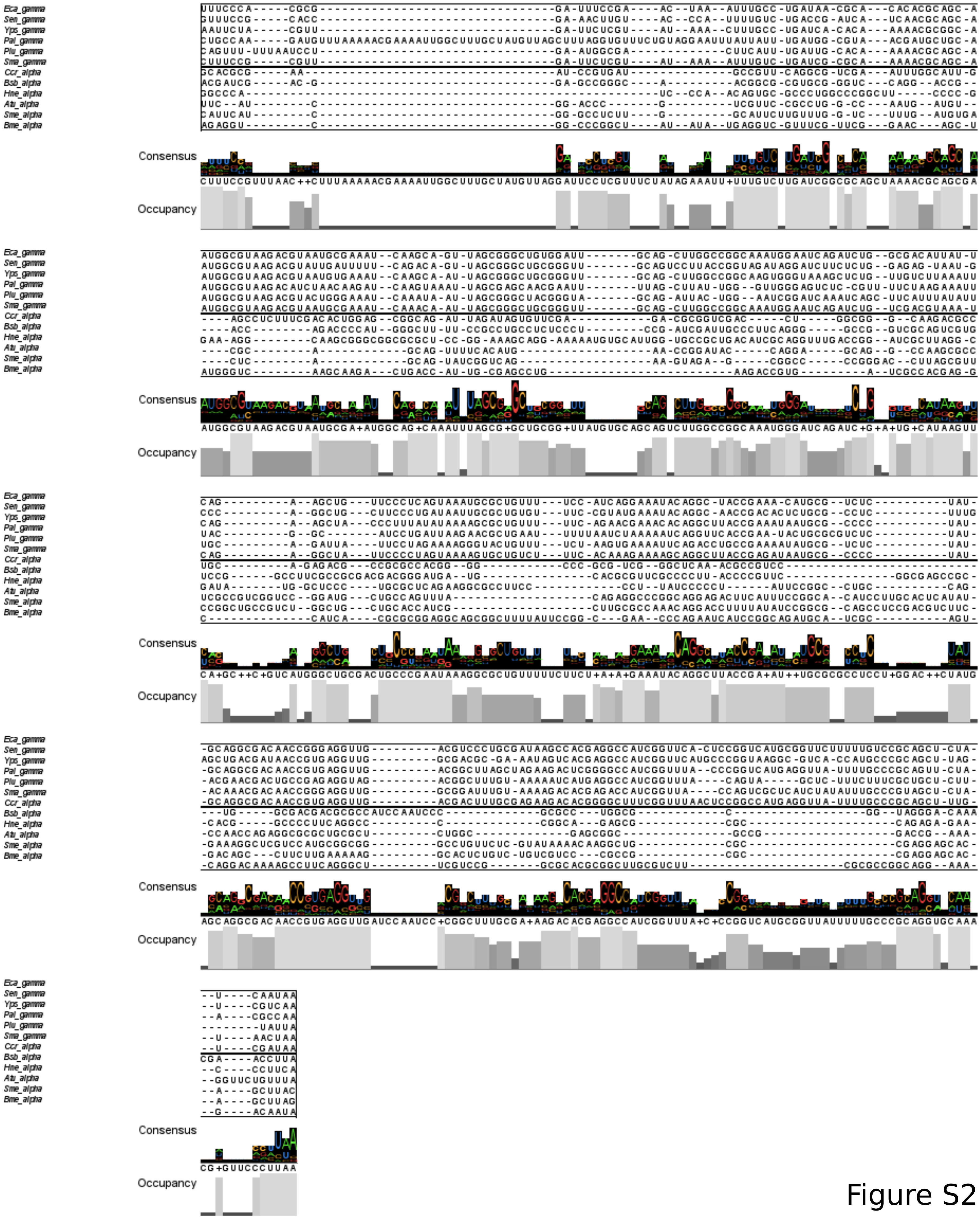
Eca_gamma
Sen_gamma
Yps_gamma
Pal_gamma
Plu_gamma
Sma_gamma
Ccr_alpha
Bsb_alpha
Hne_alpha
Atu_alpha
Sme_alpha
Bme_alpha
Consensus
Occupancy

Figure S2

### Figure S3

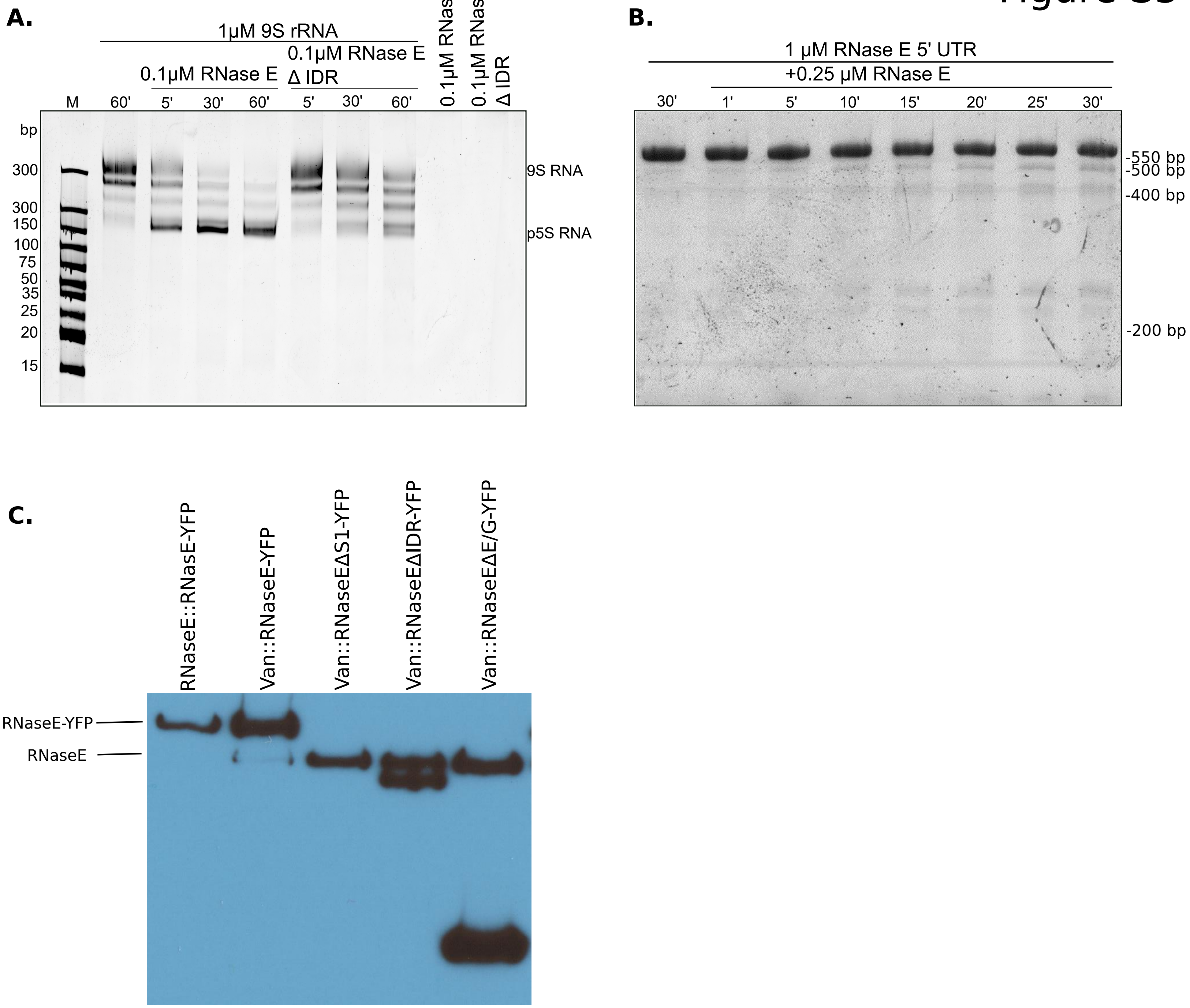


Figure S3

**Figure S1.** RNase E overexpression time course. Average YFP intensities were measured using microbe (43).

**Figure S2.** RNase E 5′ leader region sequence alignments for γ and α-proteobacteria. Sequences of RNase E 5′ leader were extracted from the genomes of six representative species of both γ and α-proteobacteria. Species names are presented using the upper case first letter representing the genus name, and the two lower case letters representing the first two letters of the species name. These sequences were aligned in turbofold (18).

**Figure S3.** RNase E domain truncations cannot autoregulate. A.) Processing of 9S ribosomal RNA (9S rRNA) to precursor 5S ribosomal RNA (p5S rRNA). Both RNase E and RNase E Δ IDR processed 9S rRNA (299 bases) to p5S rRNA (~150 bases). B.) Cleavage of RNase E 5′ UTR by RNase E. RNase E 5′ UTR was labelled at the 3′ end by Cy5. The samples were resolved on 7% Urea-acrylamide gel. RNase E variants lacking the S1 domain (JS62), the IDR (JS221), or the E/G domain (JS61) were expressed from the *vanA* locus with 0.5mM vanillate, and the probed using anti-RNase E antibody. This antibody recognizes the S1 domain of RNase E, which is why the ΔS1 sample does not show any expression. We independently confirmed ΔS1 expression by fluorescence microscopy (13).
